# Multiplexed strain phenotyping defines consequences of genetic diversity in *Mycobacterium tuberculosis* for infection and vaccination outcomes

**DOI:** 10.1101/2022.01.23.477410

**Authors:** Allison F. Carey, Xin Wang, Nico Cicchetti, Caitlin N. Spaulding, Qingyun Liu, Forrest Hopkins, Jessica Brown, Jaimie Sixsmith, Thomas R. Ioerger, Sarah M. Fortune

**Author notes:** **Correspondence:** Allison F. Carey and Sarah M. Fortune.

## Abstract

There is growing evidence that genetic diversity in *Mycobacterium tuberculosis* (Mtb), the causative agent of tuberculosis, contributes to the outcomes of infection and public health interventions, such as vaccination. Epidemiological studies suggest that among the phylogeographic lineages of Mtb, strains belonging to Lineage 2 (L2) are associated with concerning clinical features including hypervirulence, treatment failure, and vaccine escape. The global expansion and increasing prevalence of L2 has been attributed to the selective advantage conferred by these characteristics, yet confounding host and environmental factors make it difficult to identify the bacterial determinants driving these associations in human studies. Here, we developed a molecular barcoding strategy to facilitate high-throughput, experimental phenotyping of Mtb clinical isolates. This approach allowed us to characterize growth dynamics for a panel of genetically diverse Mtb strains during infection and after vaccination in the mouse model. We found that L2 strains exhibit distinct growth dynamics *in vivo* and are resistant to the immune protection conferred by Bacillus Calmette-Guerin (BCG) vaccination. The latter finding corroborates epidemiological observations and demonstrates that mycobacterial features contribute to vaccine efficacy. To investigate the genetic and biological basis of L2 strains’ distinctive phenotypes, we performed variant analysis, transcriptional studies, and genome-wide transposon sequencing. We identified functional genetic changes across multiple stress- and host-response pathways in a representative L2 strain that are associated with variants in regulatory genes. These adaptive changes may underlie the distinct clinical characteristics and epidemiological success of this lineage.

## INTRODUCTION

Pathogen population diversity can affect a range of clinically relevant phenotypes including virulence, response to treatment, emergence of antibiotic resistance, and vaccine efficacy. In order to translate a basic understanding of pathogen biology into clinical advances and begin to move towards the goal of personalized medicine in infectious diseases, it is critical to assess the generalizability of a given observation to clinical pathogen populations. With the revolution in genome sequencing, we are able to envision a future in which the features of the pathogen are incorporated into medical decision making. Rapid, inexpensive sequencing technologies have transformed our ability to enumerate the genetic diversity within and between pathogen populations. Uncovering the consequences of these genetic variants for pathogen physiology and associating them with specific phenotypes has been most successful in the arena of antimicrobial resistance. This has been possible because drug resistance can be readily and reproducibly measured *in vitro*, and there are now widely-used diagnostic assays that leverage the resulting genotype-phenotype associations to rapidly tailor antimicrobial regimens^1^. However, many clinically relevant phenotypes, such as virulence, transmissibility, or likelihood of causing different disease manifestations are less easily measured and may be confounded by variation in host features. In addition, we lack efficient experimental approaches to assess the functional consequences of pathogen genetic variation at scale and thus are limited in our capacity to create robust genotype-phenotype maps.

These challenges are particularly acute in the study of *Mycobacterium tuberculosis* (Mtb), the etiologic agent of tuberculosis, which is a leading cause of infectious disease deaths worldwide^2^. Mtb causes approximately 10 million active infections per year, and is estimated to latently infect 1/4 of the world’s population^2^. Whole genome sequencing-based phylogenetic studies have demonstrated that Mtb strains segregate into seven distinct genetic lineages (Lineages 1-7) that have geographic origins reflecting evolution concurrent with early human migration^3,4^. Epidemiological studies have found associations between strain lineage and a range of clinical phenotypes including disease progression, transmissibility, likelihood of antibiotic resistance and the efficacy of vaccination^5–13^. However, these associations are not always consistent from study-to-study^14^ and are confounded by the strong geographic structure of the Mtb phylogeny, making the impact of pathogen variation difficult to distinguish from host and health system variation^16^. Moreover, because manipulating Mtb is so cumbersome, the experimental characterization of strain differences has focused on a tiny number of reference strains, thus it is often unclear whether the identified phenotypic characteristics are reflective of lineage, sublineage, or strain level differences.

Several epidemiologic studies suggest that strains belonging to Lineage 2 (L2) are associated with hypervirulence, increased transmissibility, treatment failure, and escape from the protection conferred by vaccination^5–13^. Comparative phenotyping of an L4 reference strain, H37Rv, with an L2 strain (HN878), demonstrated that L2 strains synthesize phenolic glycolipid, a cell envelope lipid with immunomodulatory properties^15,16^, and that the associated polyketide synthase gene, *pks1/15*, is disrupted by a small deletion in L4 strains. Directed genetic studies of HN878 and H37Rv demonstrate that production of phenolic glycolipid increases virulence in mice, suggesting a model in which the increased virulence and transmission of L2 strains compared to L4 strains can be at least partially attributed to this genetic difference. However, the presence of an intact *pks15/1* open reading frame does not strictly correlate with virulence across clinical isolates. Both L2 strains and strains from the less epidemiologically successful Lineages 1 and 3, which are not associated with enhanced virulence, possess an intact *pks15/1* gene^17–19^.

The basis of other lineage-associated traits is even less well understood. L2 strains are associated with the more frequent acquisition of multidrug resistance and treatment failure and some L2 strains have an increased basal mutation rate, leading to the hypothesis that there has been selection for the evolution of hypermutability to increase fitness in the setting of widespread antibiotic treatment^20,21^. These differences in mutability have been ascribed to L2-specific missense mutations in the DNA damage repair genes *mutT2, mutT4*, and *ogt*^22,23^.

However, these variants have not been conclusively linked to hypermutability in experimental or observational studies^24–27^. L2 strains also possess genetic variants that result in the constitutive overexpression of the DosR regulon, a hypoxic response regulon hypothesized to confer a fitness advantage *in vivo* ^26–28^. However, DosR overexpression did not enhance Mtb fitness in an animal model of infection^28^. Taken together, these data suggest that it may be too simplistic to imagine that the complex clinical traits ascribed to different Mtb lineages are the result of any single mutation. Rather, the evolution of Mtb over time may have produced a network of interacting genetic variants resulting in the rewiring of key features of pathogen biology in a way that has modulated clinical characteristics. Consistent with this idea, a population genetic analysis of Mtb isolates found that non-synonymous SNPs were overrepresented in transcriptional regulators in L2 strains, a signature of selection and a potential mechanism for widespread functional genetic changes^29^.

Ultimately, to incorporate bacterial features into the design and deployment of new diagnostics and treatments for Mtb–and in infectious diseases more generally—we need facile tools to rapidly phenotype clinical pathogen populations and to define the major molecular axes of biologic variation for traits beyond antimicrobial resistance. To address these limitations for Mtb, we demonstrate the feasibility of utilizing a coordinated set of functional genomic tools to define lineage-and strain-specific virulence characteristics and map their molecular basis. We show that L2 strains exhibit broad rewiring of stress response pathways associated with variants in key regulatory genes. These adaptations may underlie this lineage’s unique clinical characteristics and global epidemiological success, and reveals vulnerabilities that could be exploited to develop improved therapeutics and more effective vaccines.

## RESULTS

### Molecular barcoding of *Mtb* clinical isolates permits multiplexed phenotyping *in vitro* and *in vivo*

We sought to develop methodology to facilitate quantitatively robust, facile phenotyping of Mtb strains. We previously demonstrated the utility of genetic barcoding to tag individual bacteria and isogenic strains in a population, which can then be assayed in experiments where competitive fitness is tracked through deep sequencing^30^. We therefore prototyped a similar strategy to rapidly define the *in vivo* characteristics of a panel of Mtb clinical isolates. We assembled a panel of 14 clinical isolates, representing three epidemiologically prevalent lineages (L2, L3, and L4), and the widely-used reference strains H37Rv and Erdman, which belong to L4 (Figure 1A)^31,32^. We tagged each strain with a unique, 8-basepair barcode that can be read out by next-generation amplicon sequencing (Figure 1B). To provide an internal assessment of experimental reproducibility, each strain was barcoded in duplicate.

**Figure 1.**
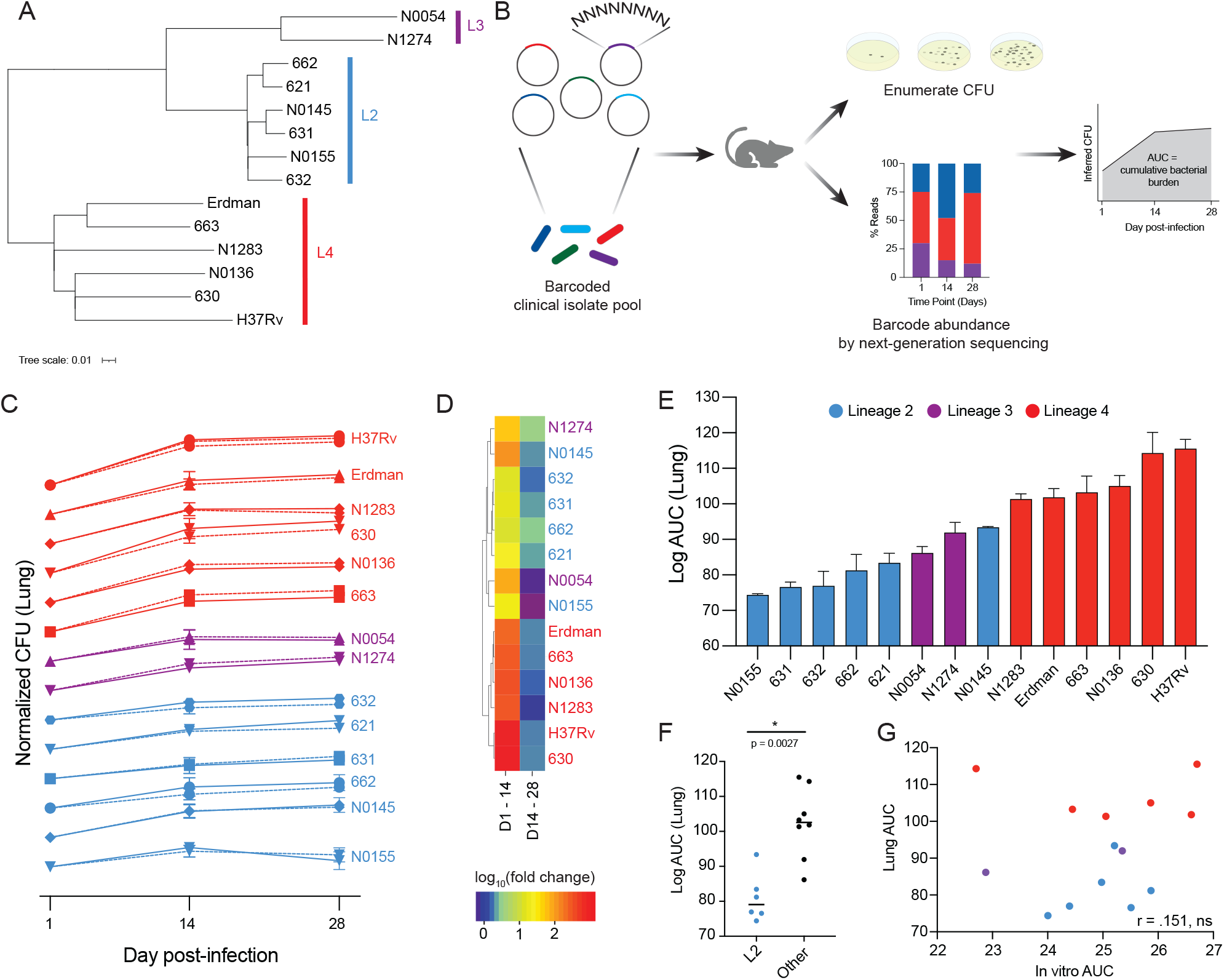
A barcoded pool of *M. tuberculosis* clinical isolates for multiplexed *in vivo* phenotyping. (A) Phylogenetic tree of *M. tuberculosis* isolates used in this study; an approximate maximum likelihood tree was generated with FastTree. (B) Strategy for barcoding and pooling isolates, performing mouse infections, calculating CFU, and determining cumulative bacterial growth. (C) Growth dynamics of *M. tuberculosis* isolates in the lung over the course of infection. Each strain’s CFU values were normalized to day 1 post-infection. Data represent means with SD (n=4). Barcode replicates are shown as solid/dashed lines. (D) Hierarchical cluster analysis of strain growth rates over the first two weeks of infection and the second two weeks of infection. (E) Cumulative growth of each strain in the lung over the four week infection. Data represent mean of replicate barcodes for each strain and SEM. (F) Growth in the lung of L2 strains compared to all other strains, significance determined by Mann-Whitney U. (G) Correlation between cumulative bacterial growth *in vitro* and *in vivo* in the lung (Pearson correlation coefficient of log_10_ transformed data).

We then evaluated the viability of this approach to enumerate strain fitness *in vitro* and in an infection model. To measure *in vitro* growth dynamics, barcoded strains were pooled and inoculated into standard media. Bacteria were plated for CFU enumeration and genomic DNA extraction on days zero, three, and seven post-inoculation. Barcode abundance was determined by amplicon sequencing (see Materials & Methods), and an inferred CFU for each strain was calculated from the total CFU and relative barcode abundance at each time point (Figure 1B). Inferred CFU values were normalized to input values. We found that growth rates of barcode replicates for each strain were highly correlated within experiments and across independent experiments (Supplemental Figure 1A, 1B, Supplemental Table 1).

Having demonstrated the capacity of this approach to robustly track bacterial strain growth dynamics *in vitro*, the barcoded pool was then used to infect C57BL/6 mice. One, 14, and 28 days post-infection, mice were sacrificed and spleen and lung tissue harvested for CFU enumeration and barcode abundance as described above. Each strain’s inferred CFU values were normalized to day one values. We found that growth rates of strain barcode replicates were highly correlated in both lung and spleen tissue (Figure 1C, Supplemental Figure 1E, Supplemental Table 1). We performed a second infection and found that strain growth rates in two independent experiments were also highly correlated (Supplemental Figure 1F). These results demonstrate that our barcoding approach permits highly reproducible, multiplexed tracking of Mtb growth dynamics over the course of infection.

### Barcoding reveals lineage-specific growth dynamics during infection

Bacterial growth *in vivo* is an essential component of pathogenicity, and different growth rates may be advantageous during different disease stages and states. Mtb growth dynamics are characterized by an initial phase of relatively unchecked growth before an effective immune response can be mounted^33^. This is followed by an extended, sometimes life-long, period of reduced bacterial burden which represents the outcome of a dynamic interplay between pathogen growth and host-mediated killing^33^. Some, but not all, animal studies have observed an increased bacterial burden among L2 strains during acute infection, a trait that is suggested to provide a selective advantage^34–36^.

Therefore, we sought to define strain and lineage growth dynamics during infection with our barcoding approach. Because the lung is the physiological niche to which Mtb is adapted, we focused on bacterial growth phenotypes in this tissue. In the lung, we observed variable growth dynamics that appeared similar among strains of the same lineage (Figure 1C). Hierarchical cluster analysis of growth rates confirmed that strains belonging to L2 grouped together, while strains belonging to L4 grouped together (Figure 1D). The growth dynamics of L4 were characterized by rapid growth over the first two weeks of infection, followed by a plateau over the second two weeks of infection. L2 growth dynamics were characterized by slower growth over the first two weeks of infection and continued, steady growth over the following two weeks (Figure 1C, 1D). Strains from L3 exhibited mixed growth dynamics.

We next assessed cumulative bacterial growth over the course of the infection by calculating the area under the curve (AUC) of the log-transformed, normalized CFU values (Figure 1B). Unexpectedly, we found that bacterial growth in the lungs over the 4-week infection period was significantly less in the L2 strains compared to other strains (p = 0.0027) (Figure 1E, 1F). Analysis of the spleen CFU data did not reveal differences in L2 cumulative growth, indicating that strain replication dynamics are tissue-specific, consistent with previous studies^37^ (Supplemental Figures 1G, 1H). L2 strains did not exhibit reduced growth *in vitro* in 7H9, a standard culture media (Supplemental Figure 1C), and there was no correlation between cumulative bacterial growth under this *in vitro* condition and *in vivo* growth (Supplemental Figure 1D), suggesting that strain growth dynamics are sculpted by the infectious environment.

### BCG confers less protection against infection by L2 strains

The L2 growth characteristics were surprising given our assumption that increased epidemiologic fitness would correlate with increased bacterial burden *in vivo*. However, more nuanced models for the increasing prevalence of L2 suggest that this lineage has become epidemiologically dominant in the setting of widespread vaccination with Bacillus Calmette-Guerin (BCG)^38^. BCG is a live, attenuated strain of *Mycobacterium bovis* whose protective efficacy is both incomplete and variable^39^. One contribution to the variable efficacy of BCG is thought to be Mtb strain diversity, and some, but not all, epidemiological studies have found that BCG has reduced efficacy against infection by L2 strains^8,36,40–43^. However, this has been difficult to assess in human population studies due to host and environmental confounders. Therefore, we next aimed to use our molecular barcoding approach to determine whether BCG confers equal protection against L2 strains compared to strains from other lineages.

Mice were vaccinated subcutaneously with BCG, rested for 12 weeks to allow an adaptive immune response to develop, then challenged with the barcoded Mtb pool (Figure 2A). One day, two weeks, and four weeks post-challenge, lung and spleen tissue were harvested, and CFU inferred as described above. To quantify protection, we calculated the difference in cumulative bacterial growth over time between naïve and BCG-vaccinated animals (Δlog_10_AUC). We found that the protection conferred by BCG vaccination varied by strain (Figure 2B). Consistent with epidemiologic predictions, BCG conferred less protection against L2 strains than other strains in the pool (p = 0.0007, Figure 2C). As we observed in our analysis of growth dynamics during primary infection, the protection conferred by BCG was tissue-specific, and there was no difference in protection between L2 strains and other strains in the spleen (Supplemental Figure 2A, 2B).

**Figure 2.**
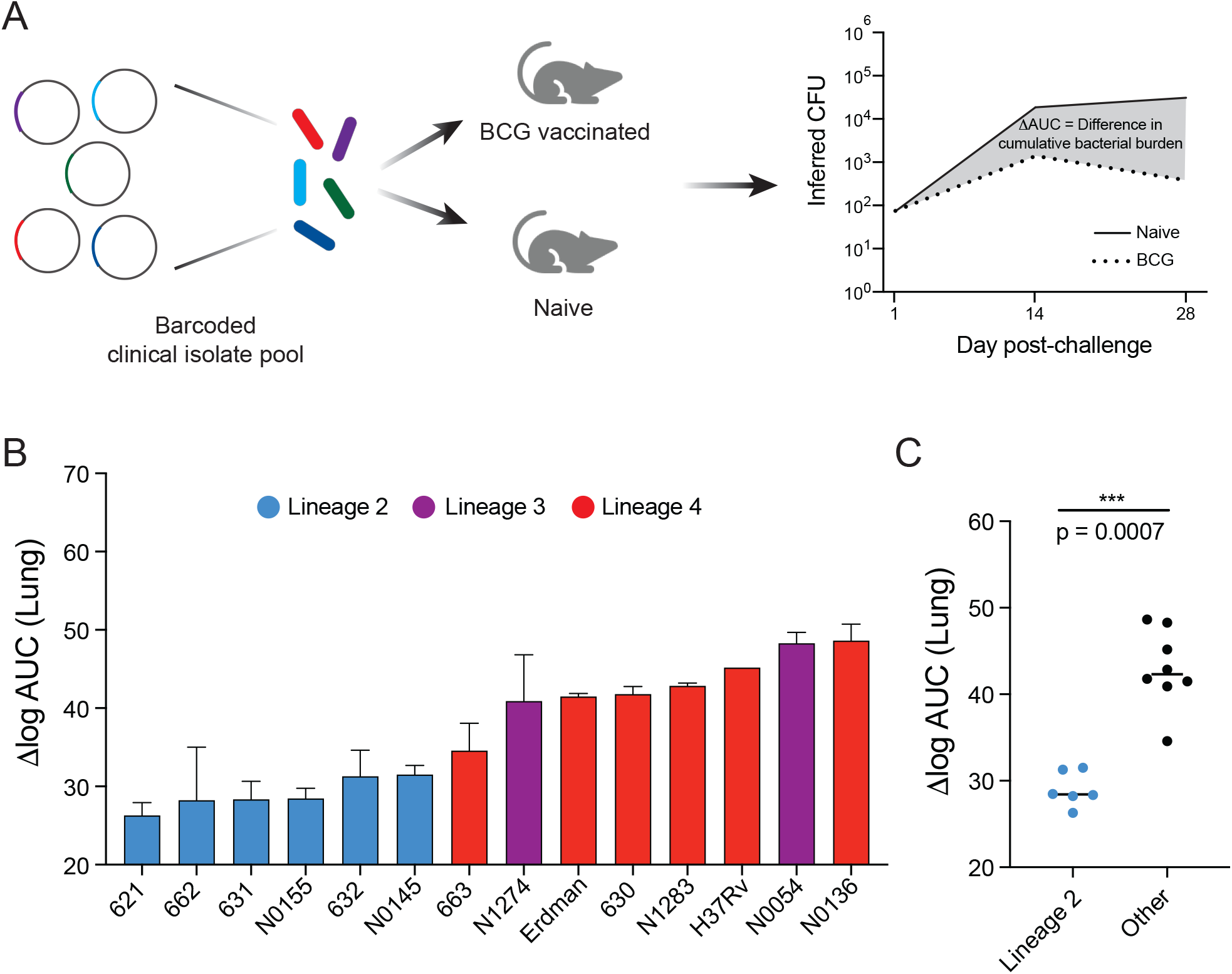
Defining strain and lineage contributions to BCG vaccine escape. (A) Strategy for vaccinating and challenging mice and quantifying protection. (B) Difference in bacterial burden in the lung conferred by BCG vaccination over the course of the four week infection. Data represent mean of replicate barcodes and SEM. (C) Protection conferred by BCG vaccination against L2 strains compared to all other strains, significance determined by Mann-Whitney U.

### Strain-specific differences in gene expression under stress conditions

Together, these data indicate that L2 strains have *in vivo* traits that are not neatly classified as “hypervirulence”. To better understand the relevance of these features to the more complex context of human infection, we sought to identify bacterial pathways shaping the *in vivo* biology of L2 strains. Comparative genomic and population genetic analyses have identified sequence variants specific to L2 strains, and found that variants in regulatory genes are overrepresented^22,29^. These genetic changes include non-synonymous SNPs in the *dosR/S/T* and *kdpD/E* two-component systems, the serine/threonine protein kinase *pknA*, the LuxR family regulators *Rv0890c* and *Rv2488c*, and the tetR family regulators *Rv0452* and *Rv0302*, among others. The impact of most of these variants for pathogenesis has not been determined, however, this sequence-level analysis suggests differential engagement of key regulatory nodes at the host-pathogen interface in L2 strains, with potential consequences for infection phenotypes.

To test this model, we selected representative L2 (621) and L4 (630) strains from the barcoded panel, in addition to the widely-used reference strain, H37Rv, which belongs to L4, for further characterization. We included a clinical isolate from L4 as a comparator because it is likely that H37Rv has adaptations due to continuous laboratory culture^44^. First, we identified genetic variants specific to L2 strain 621 compared to H37Rv and the L4 clinical isolate (Supplemental Table 2). Consistent with published studies, we identified variants in regulatory genes, including a one basepair deletion in the gene encoding the DosT sensor kinase, which has been linked to overexpression of the DosR hypoxia responsive regulon under exponential growth conditions^45^, as well as synonymous and non-synonymous SNPs in the genes encoding the MprA/B two-component system, which regulates numerous stress- and host-response pathways, including the alternative sigma factors and the ESX-1 virulence system (Figure 3A)^46,47^.

**Figure 3.**
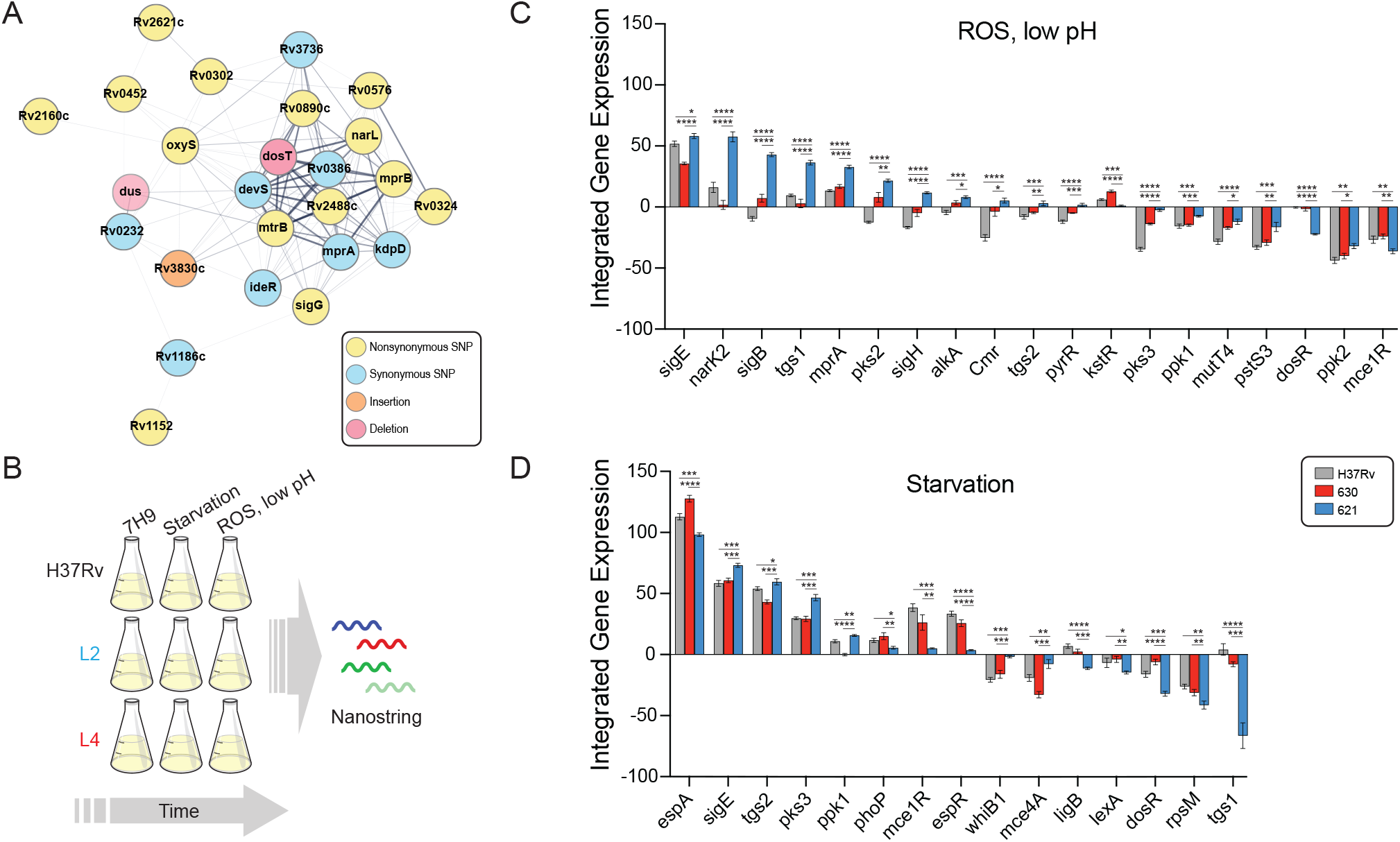
Transcriptional signatures under stress conditions differ between Mtb strains. (A) STRING plot of regulatory genes with coding region variants specifically in the L2 strain 621 as compared to the L4 strain 630 and the reference strain H37Rv. Edge thickness represents strength of evidence for direct interaction. (B) Strategy for the *in vitro* stress gene expression experiment. (C and D) Genes with quantitative and qualitative differences in expression in the L2 strain under oxidative stress, low pH conditions (C) and under starvation conditions (D) over the course of the experiment. Asterisks indicate significant differences in integrated gene expression over time determined by calculating the area under the curve for T0 normalized, log_2_ transformed data and performing one-way ANOVA with Tukey’s post-test for significance.

Given these and other genetic differences in critical regulators of bacterial adaptation to host-imposed stresses, we next assessed the transcriptional responses of these strains under *in vitro* conditions that mimic the phagolysosomal environment inhabited by Mtb, specifically, oxidative stress at low pH and nutrient starvation (Figure 3B). To do so, we designed a custom Nanostring probe set to measure expression of 54 curated bacterial stress regulators and downstream response genes (Supplemental Table 3). These targets were selected because they have been shown to be induced during infection or under *in vitro* conditions that approximate the infectious milieu^48–51^. RNA was extracted two, six, and 24-hours post stress induction and reads were normalized to internal controls and T0 (see Materials & Methods). Hierarchical cluster analysis revealed concerted changes in gene expression under each condition, consistent with prior reports (Supplemental Figure 3)^48^. Because we measured gene expression at multiple time points, we integrated normalized Nanostring counts over time for a more robust assessment of each strain’s transcriptional response. To identify L2-specific differences in expression, we filtered for genes that were both quantitatively and qualitatively differentially expressed in the L2 strain as compared to both H37Rv and the L4 clinical isolate (Figure 3C, D, Supplemental Table 4).

Among this set of differentially expressed genes, we observed higher expression of the alternative sigma factors *sigB, sigE*, and *sigH*, as well as the two-component sensor *mprA* under the low pH, oxidative stress condition (Figure 3C), and higher expression of *sigE* under starvation (Figure 3D). S*igE* is considered a master regulator of mycobacterial gene expression under stress conditions^52^, while *sigB* appears to be an end regulator in the sigma factor cascade^53^. *SigE, sigB*, and *sigH* are part of a transcriptional circuit with the MprA/B two-component system, a central sensor of environmental stresses and key determinant of mycobacterial persistence during infection^47,54–56^.

Previous studies have found that *dosR* expression is constitutively higher in L2 strains, which we also observed in the T0 data (Supplemental Table 4), however, we found that *dosR* expression was significantly lower in the L2 strain under both stress conditions (Figure 3C, 3D) ^28,45^. This suggests that the L2-specific *dosR* genetic variants alter the transcriptional response of this regulator under stress conditions as well as under basal conditions, potentially in diverging ways. A subset of the *dosR* regulon genes were included in our expression panel: *narK2* (nitrate transport), and *tgs1* (triacylglycerol synthase). Both genes were differentially expressed in the L2 strain under the tested stress conditions, displaying condition-specific expression profiles, with higher expression of *narK2* and *tgs1* under the low pH, oxidative stress condition, and decreased expression of *tgs1* under starvation. This likely reflects the integration of signals from multiple regulators to generate a response appropriate for both gene function and environmental conditions. Taken together, these targeted expression data indicate that L2 strains have a distinct transcriptional response to the stresses experienced during infection.

### Functional genomic analysis of Mtb strains during infection

An alternative to using whole-genome sequencing and expression analyses to develop models of the biological pathways driving pathogen phenotypes is instead to leverage a functional genomic method: transposon sequencing (TnSeq). TnSeq entails genome-wide transposon mutagenesis coupled with next-generation sequencing, and is a high-throughput, unbiased approach to defining bacterial genetic requirements for survival and growth under a condition of interest^57^. In contrast to sequence analyses, where the biological consequences of individual variants may be difficult to predict, or transcriptomics, which can discount the role of constitutively expressed genes and post-transcriptional regulation, TnSeq provides a functional readout of the fitness cost of gene disruption. Importantly, strain-to-strain differences in genetic requirements identified by TnSeq have been shown to reflect meaningful differences in bacterial physiology^32,58,59^. Therefore, we sought to use this approach to comprehensively define functional genetic differences in L2 strains during infection.

To do so, C57BL/6 mice were infected with saturated transposon libraries of the three strains subjected to sequence and expression analysis: L2 strain 621, L4 strain 630, and reference strain H37Rv. Because we observed the greatest differences in bacterial growth dynamics between L2 and other strains two weeks post-infection (Figure 1D), we chose one- and two-week timepoints for TnSeq analysis. TnSeq data is frequently applied to dichotomously define genes as essential or non-essential for growth under a given condition. However, a limitation of a binary classification system is that quantitative differences in genetic requirements are not uncovered. For example, a gene might be classified as non-essential in all strains, yet the relative fitness cost of disrupting the gene may differ and can reflect important physiological differences among strains^32,60^. Capturing such quantitative differences from a conditional TnSeq dataset requires accounting for differences in the input libraries that exist due to both the stochastic nature of transposon mutagenesis and biological differences among strains. To accomplish this, we applied a Bayesian method that performs a four-way comparison of transposon-junction read counts across input and output libraries, and compares the relative change in transposon mutant abundance (Figure 4A)^61^. This interaction analysis identifies genes that are conditionally essential *in vivo* in a strain-dependent manner. This pipeline was originally developed to identify epistatic genetic interactions between deletion strain and wild-type backgrounds, however, we reasoned that it could be used to identify differences in genetic requirements between strains of distinct genetic backgrounds.

**Figure 4.**
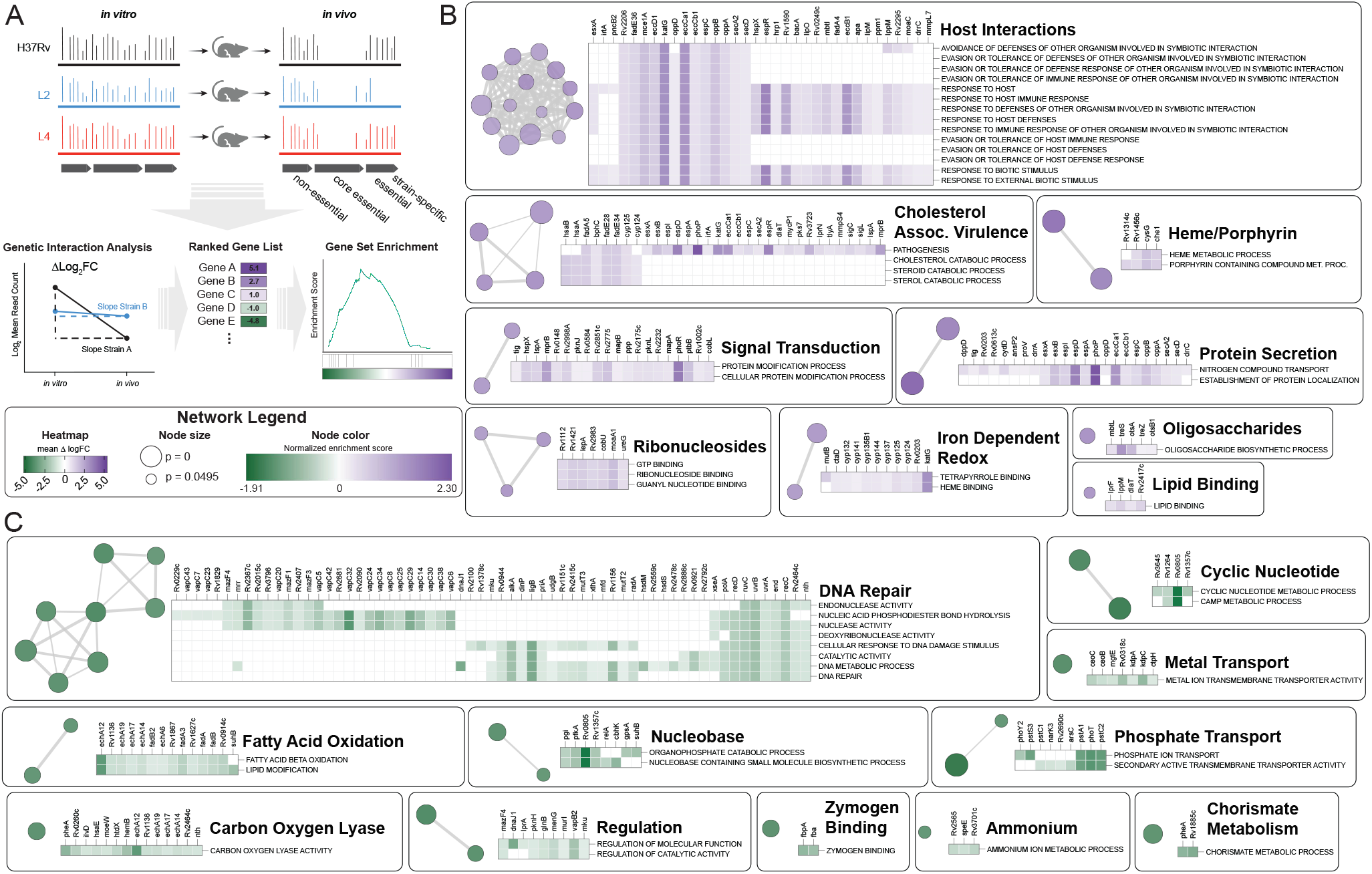
Functional genomics to identify genetic determinants of L2 infection phenotypes. (A) Experimental strategy and analytic approach to defining differences in relative genetic requirements between strains during infection using transposon sequencing and genetic interaction analysis. (B and C) Network plots generated in Cytoscape depicting genes that have a decreased requirement (B) in the L2 strain compared to the reference strain, H37Rv, one week post-infection or an increased requirement (C) by GSEA. Nodes represent enriched Gene Ontology (GO) Terms with a cutoff of p <0.05. GO Terms that were also significant in the comparison between H37Rv and the L4 clinical isolate 630 were excluded. Node color represents normalized enrichment score. Node size is inversely proportional to significance value. Edge thickness represents the number of overlapping genes, determined by the similarity coefficient. Heatmaps display leading edge genes for each cluster, with color corresponding to the Δlog_2_(fold-change) values of the genetic interaction TnSeq analysis.

We therefore performed pairwise interaction analysis between the reference strain H37Rv and each of the clinical isolates at each time point (Supplemental Table 5). To define 621-specific differences in the genetic requirements for infection, we considered only genes that were statistically significant (adj. p-value <0.05) in the H37Rv-621 comparison but not significant in the H37Rv-630 comparison. By these criteria, 32 genes were differentially required in the L2 strain one week post-infection, and 118 genes were differentially required two weeks post-infection. These gene sets were highly overlapping, as 21 of the 32 genes significant at week one were significant two weeks post-infection. To gain insight into the biological processes that differ among strains during infection, we performed gene set enrichment analysis (GSEA) on the output of the interaction analysis, using the Δlog_2_(fold-change) values as input for the preranked method and Gene Ontology (GO) Terms for functional annotation (Figure 4A)^62^. GSEA found that compared to the reference strain H37Rv, the L2 isolate had 73 significantly enriched GO Terms (p<0.05). To identify pathways that were enriched specifically in the L2 strain, 25 GO Terms that were significant in the comparison between H37Rv and 630 were excluded. The remaining 48 GO Terms indicated a decreased requirement in the L2 strain for genes involved in host interactions, including the canonical virulence system, ESX-1; cholesterol catabolism; protein secretion; and heme metabolism (Figure 4B, Supplemental Table 6). There was an increased requirement in the L2 strain for genes involved in DNA damage repair; phosphate uptake; fatty acid oxidation; and cyclic nucleotide signaling, among others (Figure 4C). We found similar differences in GO Term enrichment when comparing the L2 and L4 clinical isolates head-to-head (Supplemental Table 6), indicating that the observed differences do not simply reflect laboratory adaptation of H37Rv. Most of these processes were also enriched at the two-week time point (Supplemental Figure 4, Supplemental Figure 5), suggesting sustained, strain-specific differences in host-pathogen interactions during infection.

To place the variability in genetic requirements we observed between bacterial isolates from different phylogenetic lineages into broader biological context, we considered a recently published TnSeq study which investigated Mtb requirements for infection across genetically and immunologically diverse mouse backgrounds^63^. In this study, an H37Rv transposon library was used to infect a panel of 60 mouse genotypes encompassing strains from the Collaborative Cross collection and mice with specific immunological deficits, such as IFNγ knockout. This approach facilitated a comprehensive assessment of variation in bacterial genetic requirements under distinct infection conditions. Consistent with our work and previous studies, the authors identified 234 genes required for H37Rv to grow or survive in C57BL/6 mice, yet there were as many as 212 additional *in vivo*-essential genes per mouse genotype. This is comparable to the 172 genes we identified as differentially required to infect C57BL/6 mice in the L2 isolate compared to H37Rv, suggesting that the functional genetic differences between Mtb strains can be as substantial as those that are imposed by distinct host backgrounds. Through network analysis, the authors found that differentially required genes could be clustered into 20 modules with correlated changes in fitness. We performed a statistical analysis of the overlap between these modules and the genes that were differentially required in the L2 strain during infection and identified three modules with significant overlap (p-adj <0.05, Fisher’s exact test). These modules are categorized as ESX-1, phosphate uptake, and an uncategorized set that includes a number of DNA damage repair genes. This intersection of host- and pathogen variability suggests that certain lineages of Mtb may be adapted to specific host environments, consistent with population genomic analyses^4^.

### Regulatory variants are associated with differential genetic requirements during infection

Our TnSeq data indicate widespread functional genetic differences between Mtb strains over the course of infection. We noticed that many of the GO Terms found to be enriched by GSEA in the L2 strain represent biological processes regulated by genes with 621-specific genetic variants. For example, cholesterol metabolism genes are differentially required in 621, and this strain possesses a SNP upstream of *kstR*, which controls the cholesterol catabolism regulon. This suggests that rewiring of the bacterial response to the host environment may be driven by selection on regulatory genes, consistent with sequence analyses of L2 genomes^22,29^.

To test this hypothesis, we mined a published data set from a comprehensive Mtb transcription factor overexpression (TFOE) study^64^. In this work, 206 of the 214 known and predicted Mtb transcription factors were inducibly overexpressed and transcriptional signatures assessed by high-density microarray, reflecting both direct and indirect regulatory effects. We integrated this data with our TnSeq results to determine which differentially required genes (as determined by genetic interaction analysis) were regulated by transcription factors with sequence variants. In cases such as the DosR regulon, where variants are located in the sensor of a two-component system, we considered genes regulated by the transcription factor. We found that 42 of the 129 genes that were differentially required specifically in L2 strain 621 were regulated by a transcription factor possessing a 621-specific genetic variant (Figure 5A). To assess the statistical significance of this finding, we performed a simulation with a null distribution of 10,000 trials of 129 genes chosen at random and found the overlap to be highly significant (p = 0.0048). This result is consistent with a model in which variants in response regulators drive functional genetic differences among strains.

**Figure 5.**
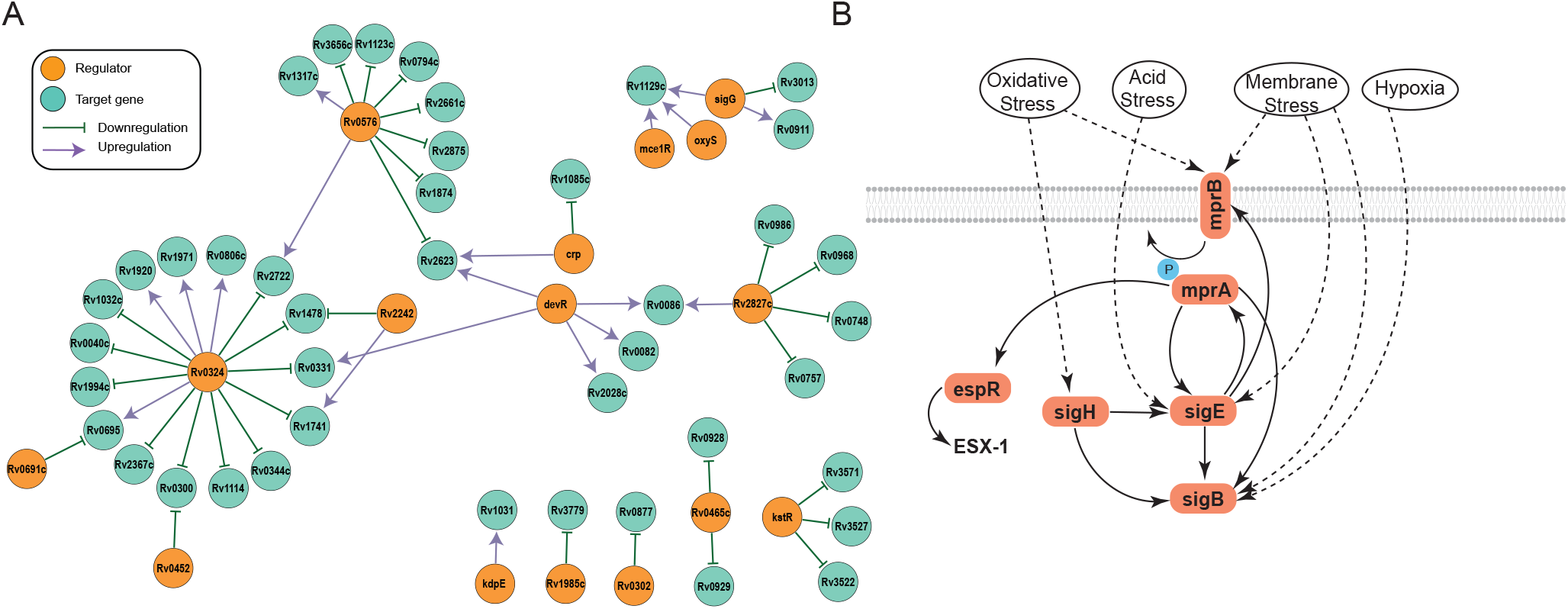
Differentially required genes are regulated by transcription factors with strain-specific variants. (A) Network plot generated in Cytoscape showing genes with L2-specific TnSeq differences that are transcriptionally regulated by systems with strain-specific genetic variants. (B) Schematic depicting the complex regulatory circuit of the two component system MprA/B, which has a nsSNP in the sensor gene *mprB* in strain 621.

This analysis likely underestimates the relationship between genetic variants in transcriptional regulators and the differential genetic requirements identified by TnSeq in this representative L2 strain. In the TFOE study, transcriptional responses were assessed under a single, *in vitro* growth condition at a single time point, and stringent statistical thresholds were used to determine regulatory relationships. This may mask subtle but biologically important regulatory roles. For example, the transcriptional activator *mprA*, part of the MprA/B two component system, was not found to regulate any genes by the rigorous thresholds of the TFOE study. However, directed genetic studies have found that *espR* is regulated by *mprA*^46,65^, and the sensor kinase of this system, *mprB*, has a non-synonymous SNP in strain 621 (Figure 5B). *EspR* regulates the ESX-1 virulence system, which was differentially required by the L2 strain during infection (Figure 4B, Supplemental Table 5). MprA/B is also part of a regulatory loop with the alternative sigma factors *sigB, sigE*, and *sigH*, therefore, genetic variants at the top of this cascade may have pleotropic transcriptional effects.

## DISCUSSION

Tuberculosis is a notoriously heterogeneous disease, with outcomes ranging from lifelong, symptomatic latency to primary progressive disease. Dissecting the impact of bacterial genetic variation to this heterogeneity has been limited by confounding host and environmental factors in population studies, and by the experimental intractability of Mtb in laboratory studies. Here, we developed a robust molecular barcoding approach that allowed us to characterize *in vivo* growth dynamics in a high-throughput fashion for a genetically diverse panel of isolates. Among these isolates are strains from L2, a lineage that has been expanding in population size over the past two centuries, possibly due to traits that confer a selective advantage^66^. One of the features attributed to L2 strains in some epidemiological and small animal studies is increased virulence^6,8,9,13,16^. Therefore, it was unexpected that our *in vivo* fitness phenotyping revealed reduced cumulative bacterial growth of L2 strains over the course of infection compared to other strains. An explanation for this discrepancy may be that many previous animal studies used a single strain or small number of strains isolated from outbreaks, such as HN878, which might inadvertently bias towards hypervirulence. In this study, we included L2 strains from a reference set that was curated to be representative of each lineage^31^. Thus, slower bacterial growth during acute infection may be more typical of the growth dynamics of L2 than prior studies suggested. Indeed, Mtb is a pathogen that can infect an individual for a lifetime without a measurable increase in bacterial burden, and slow growth may be a survival strategy that circumvents immune-mediated killing^67^. Therefore, perhaps it is not surprising that an epidemiologically successful lineage of Mtb exhibits reduced growth compared to other strains, at least during the early stages of infection.

Our barcoding approach also permitted a systematic examination of Mtb strain and lineage contributions to the efficacy of BCG vaccination, an unresolved question in the field. The importance of Mtb strain variation for vaccine efficacy have been difficult to assess in population studies, where host and environmental factors also vary. Our findings in the mouse, a relevant pre-clinical model for Mtb vaccination studies, experimentally confirm observations made in some epidemiological studies of reduced BCG efficacy against L2 strains. This suggests that as new tuberculosis vaccines are designed, they should be evaluated for efficacy against genetically diverse and epidemiologically prevalent strains, and our barcoding approach provides a scalable means to do so.

Together, these studies demonstrate the power of molecular barcoding for high-throughput phenotyping of bacterial strains, an approach that is applicable to numerous pathogens. Although only one mouse genotype was used in the infection and vaccination studies, the C57BL/6 background is widely-used and recapitulates many features of human tuberculosis^68^. Importantly, our barcoding method makes future studies in diverse host backgrounds experimentally tractable. A limitation of barcoding is that it does not permit investigations of immune-mediated disease due to the multiplexed nature of the experiments. Although robust measurements of bacterial growth can be performed with this method, bacterial burden is not the only feature driving virulence, and differences in immunopathology may drive differences in disease severity and transmission that we cannot capture. Another limitation is that phenotypes that trans-complement will not be uncovered, however, this is a feature of other pooled phenotyping techniques, such as TnSeq and CRISPRi, which have nevertheless revealed important biological principles about numerous pathogens. Despite these limitations, as we demonstrate here, phenotypically distinct bacterial isolates can be identified for subsequent high resolution, single-strain characterization.

Mtb is an obligate human pathogen that is exquisitely adapted to the hostile environment of the lung and has evolved a suite of mechanisms to survive the stressors it encounters during infection^69^. Our in-depth genetic, transcriptional, and functional genomic characterization of representative isolates indicate that the L2 strain is functionally rewired across many of these pathways. The genes we identified by TnSeq with L2-specific differential requirements during infection represent key adaptive processes including the ESX-1 virulence system, lipid metabolism, and DNA damage repair. Our analysis indicates that these differentially required genes are more likely to be regulated by transcription factors with strain-specific variants than chance, a potential mechanism of evolutionary adaptation. Population genomic analyses are consistent with this observation, having found that transcriptional regulators are enriched for variants in L2^22,29^. Indeed, studies across other prokaryotic species suggest that evolution of transcription factor network structure is an important means of phylogenetic diversification and can lead to the emergence of organisms with distinct responses to environmental stimuli^70^.

A limitation of our transcriptional and functional genomic studies is that only one clinical isolate from L2 and L4 was characterized. The selected strains were representative of their lineage in growth characteristics and genetic features. However, in addition to lineage-level genetic diversity, strain-level genetic diversity has the potential to affect pathogenic traits. Variants present in some, but not all, strains within a lineage represent an evolutionary sandbox for selection, and dissecting the consequences of both levels of genetic variation for bacterial fitness can help define the selective landscape shaping Mtb’s ongoing adaptation. Such studies are now feasible with barcoding, which can facilitate phenotyping of numerous strains at-scale under a range of *in vitro* and *in vivo* conditions. Coupled with computational techniques such as bacterial genome-wide association, the pathogen genes and variants that drive infection outcomes and response to clinical interventions such as vaccination can be uncovered, leading to the development of molecular diagnostics to guide more effective clinical care.

## MATERIALS & METHODS

### Bacterial strains

Clinical strains were identified as previously described and cultured from single colonies^4,31^. Strains were grown at 37°C and cultured in Middlebrook 7H9 salts supplemented with 10% OADC, 0.5% glycerol and 0.05% Tween-80 or plated on 7H10 agar supplemented with 10% OADC, 0.5% glycerol and 0.05% Tween-80 unless otherwise noted. Clinical strains were handled to minimize *in vitro* passaging. Strains were previously whole genome sequenced as described^31,32^. To compare genomic variants between H37Rv, L2 strain 621, and L4 strain 630, a custom assembly and variant calling pipeline was used as previously described^32^.

### Animals

Female C57BL/6 mice were purchased from Jackson Laboratories (Bar Harbor, Maine). Mice were 6-8 weeks old at the start of all experiments. Infected mice were housed in BSL3 facilities under specific pathogen-free conditions at HSPH. The protocols, personnel, and animal use were approved and monitored by the Harvard University Institutional Animal Care and Use Committee. The animal facilities are AAALAC accredited.

### BCG vaccination

Bacillus Calmette-Guerin originally obtained from Statens Serum Institute was prepared as previously described^71^. Mice were immunized with 100 uL of OD600 1.0 frozen bacterial culture (2e7 CFU) subcutaneously in the left flank. Mice were rested for 12 weeks post-vaccination prior to challenge.

### Barcoded clinical isolate growth *in vitro*

Mtb strains were tagged with a random 8-basepair barcode essentially as described^30^. Single colonies of each strain were picked and Sanger sequenced to identify the barcode; colonies with two unique barcodes for each strain were selected. Barcoded strains were grown to log phase, pooled, and frozen into aliquots. An aliquot was subsequently inoculated into 7H9 media, grown to mid-log phase, then back-diluted to an OD of 0.01 in 7H9 in triplicate and incubated with shaking at 37°C. At the indicated time points, an aliquot was removed from each replicate for CFU enumeration, and an aliquot removed for plating to recover ∼5e3 CFU as estimated by OD600 of the culture. Recovered CFU were scraped for genomic DNA extraction, amplicon Illumina sequencing, and barcode abundance quantification by custom Python scripts, essentially as described^30^.

### Barcoded clinical isolate mouse infections and analysis

An aliquot of the barcoded strain pool was used for tail vein infection at 1e6 CFU/mouse. At indicated time points post-infection, spleens and lungs were harvested, homogenized, and plated on 7H10 supplemented with glycerol, Tween, OADC, and 20 mg/mL kanamycin. After 3 weeks of incubation, CFU were enumerated and 1e4 CFU were scraped for genomic DNA extraction, amplicon Illumina sequencing, and barcode abundance quantification by custom Python scripts, essentially as described^30^.

### Gene expression

For oxidative and starvation stress conditions, triplicate cultures of the indicated strains were grown to mid-log phase in 7H9, pelleted and washed once in an equal volume of TBS supplemented with 0.05% Tyloxapol, then resuspended in freshly-made stress media as detailed below, or 7H9 with 0.05% Tyloxapol. For oxidative stress, bacteria were resuspended in 7H9 with 0.05% Tyloxapol buffered to pH 4.5 with 10 μg/mL menadione. For starvation, bacteria were resuspended in TBS with 0.05% tyloxapol. Cultures were incubated at 37°C with shaking, and aliquots removed for RNA extraction at the indicated time points. RNA was isolated essentially as described and quantified by Qubit RNA Assay (Thermo Fisher)^32^. 125 ng of RNA was used as input in a Nanostring assay with a custom-designed probe set (Nanostring Technologies). Target sequences are listed in Supplemental Table 3. Data were analyzed with nSolver version 4 (Nanostring Technologies); raw Nanostring counts were normalized to internal positive controls to correct for technical variation between assays, and normalized to housekeeping genes (*ansA, aceAa, secA2*) to correct for variation in RNA input (Supplemental Table 4). Normalized counts were expressed as log_2_ (fold-change) relative to T0 and data clustering was performed in R v4.0.3 using complete linkage and Euclidean distance. For statistical comparisons between strains, AUC of the log_2_ (fold-change) expression data over time were calculated and one-way ANOVA with Tukey’s post-test performed in R v4.0.3 (Supplemental Table 4).

### Transposon library mouse infections and analysis

Mice were infected via tail vein injection with 2e6 CFU of frozen aliquots of previously generated H37Rv or clinical strain *Himar1* transposon libraries^32^. At the indicated time points post-infection, spleens were harvested, homogenized, and plated on 7H10 supplemented with glycerol, Tween, OADC, 0.2% Cas-amino acids (Difco) and 20 mg/mL kanamycin. For each mouse, 1e6 surviving colonies were scraped after 3 weeks for genomic DNA extraction and transposon-junction sequencing essentially as previously described^32^. Reads were mapped to the H37Rv genome, and statistical comparisons of read counts between conditions and strains were performed using Transit v3.2.0^72^. To identify differences in genetic requirements during infection between strains, the Transit genetic interaction (GI) method was used^61^. Repetitive regions, deleted genes, and genes in a large duplicated region in the L2 strain 621 were excluded as previously described (Supplemental Table 5)^32^. Gene-set enrichment analysis and leading edge analysis were performed on the Transit GI-generated Δlog_2_fold-change values using the GSEA v4.1.0 preranked tool^62^. Genes classified as essential for *in vitro* growth in at least two of the three isolates were excluded from GSEA (Supplemental Table 7). To identify *in vitro* genetic requirements for each strain, the Transit Hidden Markov Model (HMM) method was used, with insertions in the central 90% of each open reading frame considered, and a LOESS correction for genome positional bias^73^.

## SUPPLEMENTAL INFORMATION

**Supplemental Figure 1**

(A) Growth dynamics of barcoded *M. tuberculosis* isolates in 7H9 media. Each strain’s pseudo-CFU values were normalized to input. Data represent means with SD (n=3). Barcode replicates are shown as solid/dashed lines.

(B) Correlation between bacterial growth rates in independent *in vitro* experiments (Pearson correlation coefficient of log_10_ transformed data).

(C) Cumulative growth of each strain *in vitro* comparing L2 strains to all other strains, significance determined by Mann-Whitney U.

(D) Correlation between bacterial growth rate *in vitro* and *in vivo* in the spleen (Pearson correlation coefficient of log_10_ transformed data).

(E) Growth dynamics of *M. tuberculosis* isolates in the spleen over the course of infection. Each strain’s CFU values were normalized to day 1 post-infection. Data represent means with SD (n=4). Barcode replicates are shown as solid/dashed lines.

(F) Correlation between bacterial growth rates in the lung in independent *in vivo* experiments (Pearson correlation coefficient of log_10_ transformed data).

(G) Cumulative growth of each strain in the spleen over the four week infection. Data represent mean of replicate barcodes for each strain and SEM.

(H) Growth in the spleen of L2 strains compared to all other strains, significance determined by Mann-Whitney U.

**Supplemental Figure 2**

(A) Difference in bacterial burden in the spleen conferred by BCG vaccination over the course of the four week infection for each strain. Data represent mean of replicate barcodes and SEM.

(B) Protection conferred by BCG vaccination against L2 strains compared to other strains in the spleen. Significance determined by Mann-Whitney U.

**Supplemental Figure 3**

Heatmaps of Nanostring gene expression for H37Rv, 621, and 630 strains under oxidative stress and low pH (A) and starvation (B). Each gene’s counts were normalized to input (T0) values and expressed as log_2_(fold-change).

**Supplemental Figure 4**

Network plots generated in Cytoscape depicting GSEA of genes with differential requirements in the L2 strain compared to the reference strain, H37Rv, two weeks post-infection. Nodes represent enriched Gene Ontology (GO) Terms with a cutoff of p <0.05. GO Terms that were also significant in the comparison between H37Rv and the L4 clinical isolate 630 were excluded. Node color represents normalized enrichment score. Node size is inversely proportional to significance value. Edge thickness represents the number of overlapping genes, determined by the similarity coefficient. Heatmaps display leading edge genes for each cluster, with color corresponding to the Δlog_2_(fold-change) values of the genetic interaction TnSeq analysis.

**Supplemental Figure 5**

Line plots showing log_2_(fold-change) trajectories over the course of the two week infection for leading edge genes of selected functional groupings found to be enriched by GSEA of the TnSeq data (H37Rv v. L2 strain 621). Thin lines represent individual genes, thick lines represent the average for each functional grouping.

**Supplemental Table 1. Barcode reproducibility**. Pearson correlation coefficients for strain barcode replicates calculated from normalized, log_10_ transformed pseudo-CFU of *in vitro*, spleen, and lung data. The H37Rv correlation coefficient represents the average of the three pairwise barcode comparisons.

**Supplemental Table 2. Genetic variants in L2 strain 621**.

**Supplemental Table 3. Nanostring target sequences**. Target sequences for gene expression experiments.

**Supplemental Table 4. Nanostring gene expression data**. Nanostring counts normalized to internal control and housekeeping probes for each of the three replicates at T0 and each time point under the *in vitro* stress conditions, and average AUC values and standard error derived from log_2_(fold-change) data normalized to the T0 values for each strain. Significance determined by one-way ANOVA.

**Supplemental Table 5. TnSeq Transit genetic interactions output**. Genetic interaction analysis of pairwise comparisons between each of the three strains’ input and mouse output transposon junction sequencing data. Repetitive elements, deleted genes, *in vitro* essential genes, and genes in a large duplicated region in strain 621 were removed as appropriate for each pairwise comparison such that only genes that are intact and not duplicated in both strains were included.

**Supplemental Table 6. GSEA analysis of TnSeq data**. Output of gene set enrichment analysis using the preranked method, with the Transit genetic interactions Δlog_2_(fold-change) values used as input.

**Supplemental Table 7. TnSeq Transit HMM output**. Gene essentiality calls generated from the *in vitro* H37Rv, 621, and 630 libraries. Repetitive elements, deleted genes, and a large duplicated region in strain 621 have been removed as detailed in Materials & Methods.

## ACKNOWLEDGEMENTS

We thank all members of the Fortune lab and Dr. Eric Rubin’s lab for technical advice and thoughtful discussions, in particular, Dr. Nathan Hicks, Dr. Greg Babunovic, and Sydney Stanley; Shoko Wakabayashi for assistance with mouse experiments; Dr. Amanda Martinot for kindly providing the BCG; Dr. Sebastien Gagneux for providing the *M. tuberculosis* clinical isolates; and Mr. Larry Pipkin and Dr. Noman Siddiqi, who manage the Harvard T.H. Chan School of Public Health BSL-3 facilities.

This work was supported by National Institutes of Health grants P01 AI132130 (S.M.F.); T32 AI007061 (A.F.C.), T32 CA009216 (A.F.C.) and K08 AI139339 (A.F.C.).

## Notes

### Competing Interest Statement

The authors have declared no competing interest.

