## Supplemental Figures for "Multiplexed strain phenotyping defines consequences of genetic diversity in *Mycobacterium tuberculosis* for infection and vaccination outcomes"

Supplemental Figure 1

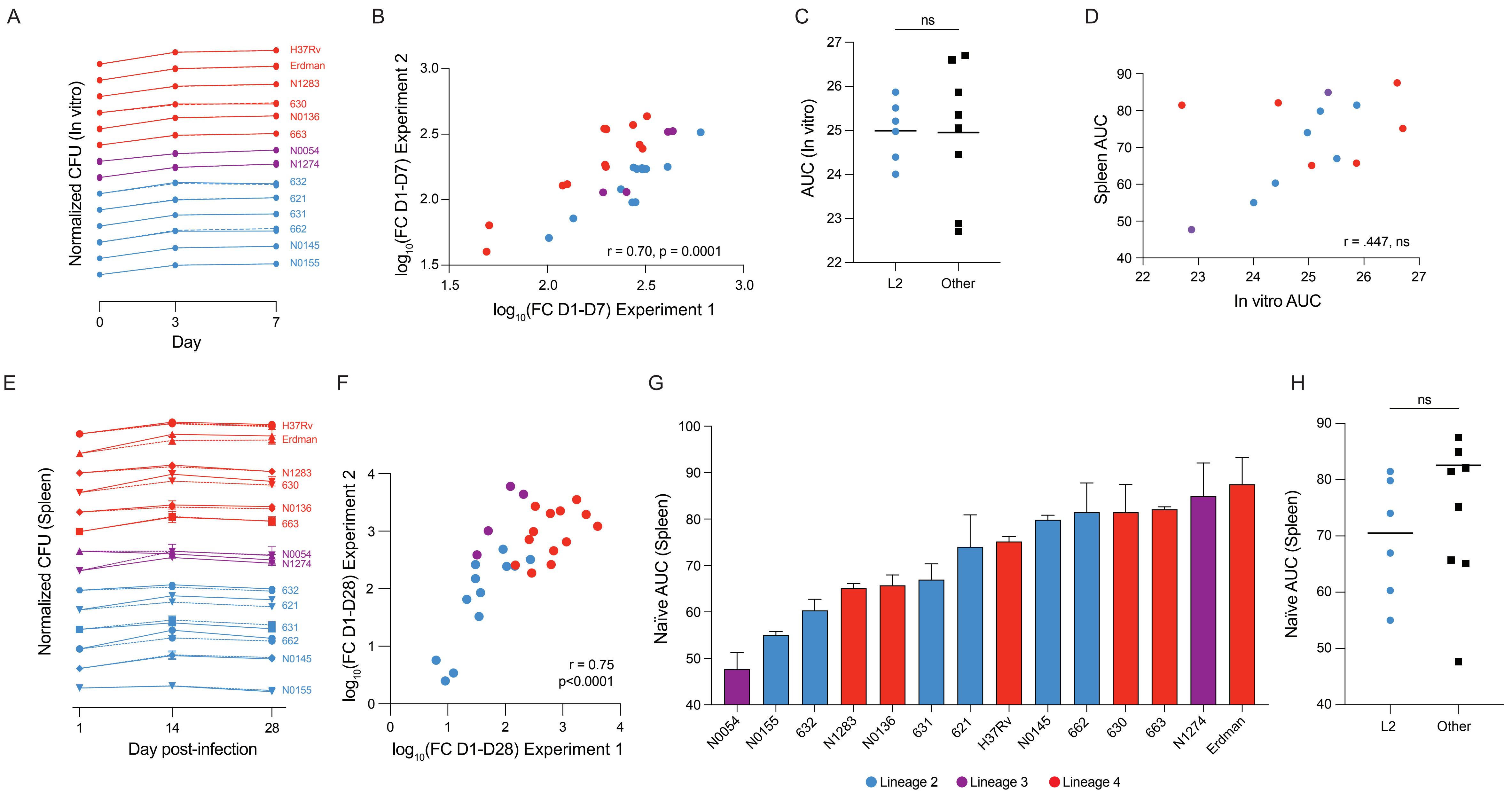

Supplemental Figure 2

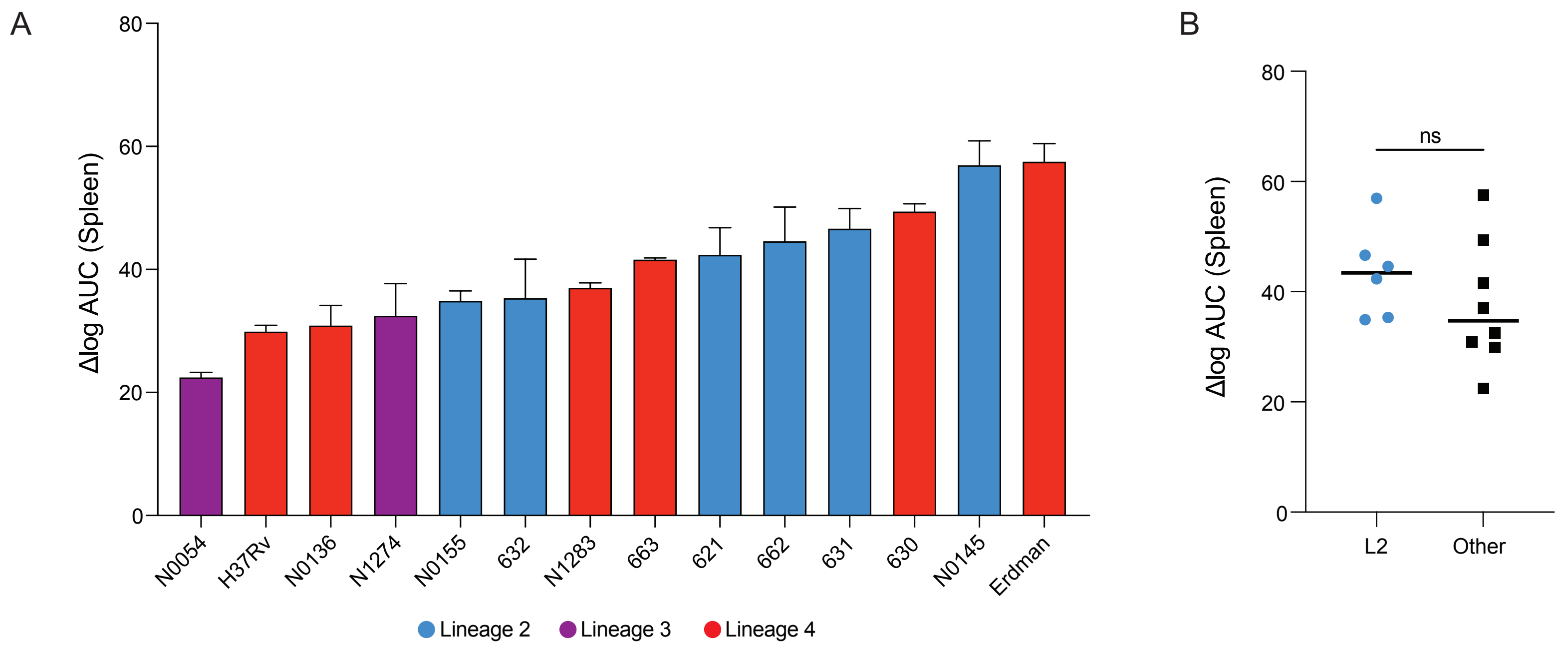

### Supplemental Figure 3

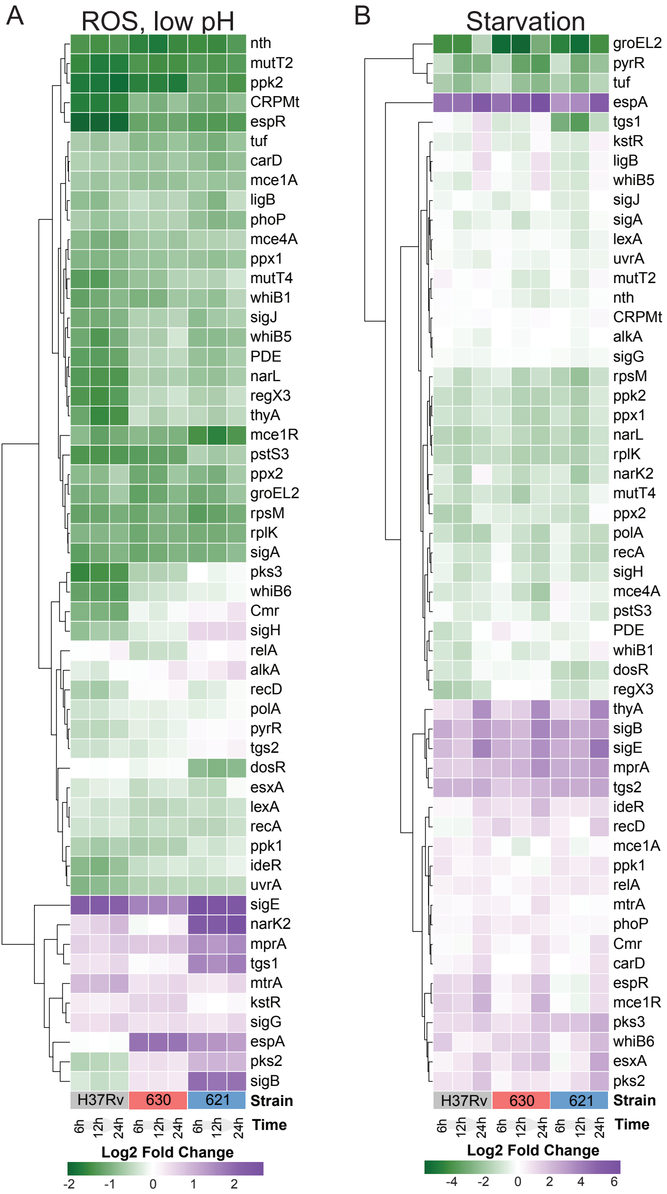

### Supplemental Figure 4

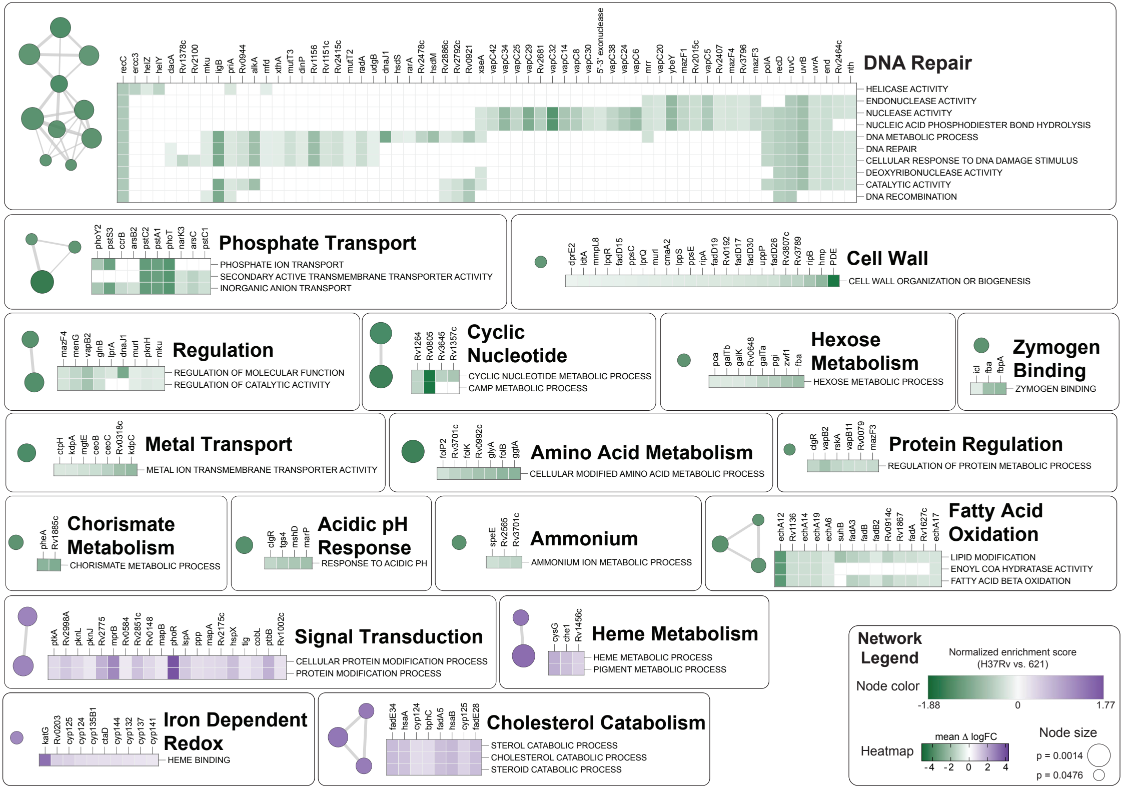

### Supplemental Figure 5

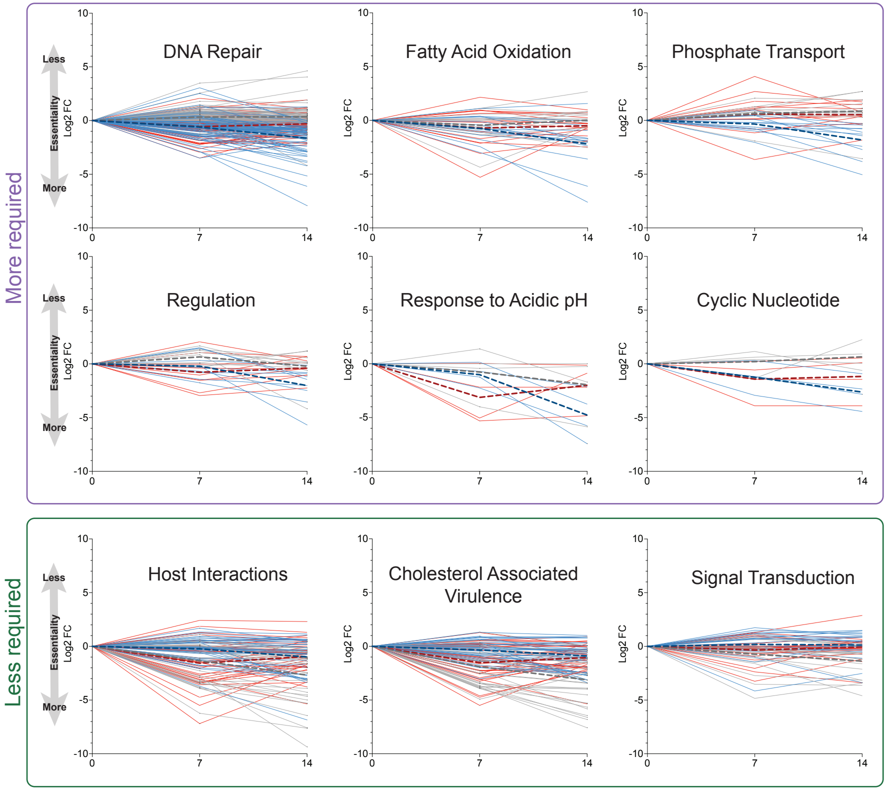
